# Genome-wide association mapping for grain and forage quality traits in a subtropical oat germplasm collection adapted to highland regions of Eastern Africa

**DOI:** 10.1101/2025.11.26.690820

**Authors:** Lidya Ashenafi, Mesfin Dejene, Kedir Mohammed, Tilahun Mekonnen, Fekede Feyisa, Hailu Lire, Alemayehu Teressa Negawo, Atikur Rahman, Susanne Barth, Jose De Vega, Chris S. Jones, Abel Teshome

**Affiliations:** Ethiopian Institute of Agricultural Research, P.O. Box 31, Holeta, Ethiopia; Addis Ababa University, Institute of Biotechnology, Addis Ababa University, P.O. Box. 1176, Addis Ababa, Ethiopia; International Livestock Research Institute P.O. Box 5689, Addis Ababa, Ethiopia; Teagasc, Crops Research Centre, Oak Park, Carlow. R93 XE12; Earlham Institute, Norwich Research Park, Norwich NR4 7UZ, UK; International Livestock Research Institute, P.O. Box 30709, Nairobi 00100, Kenya

**Author notes:** **Corresponding authors**: Teshome A.

**Keywords:** Climate-resilience, Grain and forage quality, GWAS, Oat, Population structure

## Abstract

Oat (*Avena sativa* L.) is a globally important cereal cultivated for both livestock feed and human nutrition. However, its productivity is increasingly constrained by both biotic and abiotic stresses, exacerbated by rapid climate change. To support the development of dual-usage, stress-resilient and high-yielding cultivars adapted to subtropical agroecologies, we evaluated the genetic structure and agronomic performance of a 169-member oat association panel. This panel was phenotyped for key vegetative, forage feed quality and grain related traits across three subtropical locations in Sub-Saharan Africa (SSA) over two growing seasons and genotyped using Genotyping-by-Sequencing (GBS). Diversity analysis, based on filtered SNPs, showed moderate genetic diversity (He = 0.39 and PIC = 0.3), and population structure analysis identified two main sub-groups with varying degrees of admixture, suggesting gene flow among different groups. Genome-wide association analysis uncovered 42 SNPs significantly associated with sixteen traits (false discovery rate, FDR, <0.05), along with 46 candidate genes located near these loci. Notably, candidate genes associated with seed traits included a homolog of rice *OsAK3*, encoding adenylate kinase, and a UDP-glucosyltransferase homologous to rice *GSA1,* both previously shown to regulate grain size. These genes were located near QTL linked to seed length, width, and thousand-grain weight. Another key candidate, encoding a subunit of the ESCRT-II complex (VPS25), was identified near a separate QTL and is implicated in intracellular trafficking, endosperm development, and overall seed quality. By illuminating the genetic architecture of economically important traits and identifying molecular markers linked to yield, quality and resilience, this study provides valuable genomic resources to support genomics-based breeding aimed at developing climate-resilient, productive oats varieties adapted to sub-tropical agro-ecologies.

## Introduction

Oats (*Avena sativa* L., 2n = 6x = 42) is a self-pollinated annual grass belonging to the Poaceae family and ranks as the sixth most important cereal crop globally (Rines et al., 2006; Yan et al., 2016). Valued for its versatility, oat is cultivated as a dual-purpose crop, serving both grain and forage needs. Despite its significance, the use of oats as a forage crop has received comparatively little attention in genetic research and breeding. Oat forage offers high-quality fodder and superior biomass production within a shorter growing season, making a critical feed resource during the winter season in many regions (Isidro-Sánchez et al., 2020). Although typically associated with mid-latitude climates (35° to 66.5° N), oats demonstrated broad adaptability. In subtropical areas, spring oats can be sown in autumn to avoid summer drought stress, thereby serving as a valuable source of winter forage (Esvelt Klos et al 2016). Oat is particularly well-suited to challenging environments requiring lower agricultural inputs compared to other cereal crops (Bernas et al., 2021; Omondi et al; 2022; Kamal et al., 2022). Additionally, oats contribute to sustainable agricultural practices as an effective cover crop in rotation arable systems (Tomar & Singh, 2024; Singh et al., 2025).

Given its adaptability and multifunctional role, oats present significant potential for sustainable agriculture, particularly if current production challenges can be addressed through targeted crop improvement strategies. Over recent decades, oats production has declined markedly, largely due to competition from higher-yielding cereal crops and increasing vulnerability to biotic and abiotic stresses, which are being exacerbated by effects of climate change (Kapoor & Singh, 2020; Carlson et al., 2023). Addressing these production challenges requires a nuanced approach that considers regional agro-ecological conditions and specific end-use demands. This study focuses on oat cultivation in sub-Saharan Africa (SSA), particularly in Ethiopia, where oat is predominantly grown as a winter crop. In this context, the primary breeding traits are traits related to biomass production and grain quality, aligned with its dual-purpose use.

Genetic diversity studies are essential for both conservation of genetic resources and the identification of superior genotypes for crop improvement (Arora et al., 2021; Begna, 2021). In oats, several valuable studies have explored genetic diversity using a range of molecular markers, including amplified length fragment polymorphism (ALFP) (Achleitner et al., 2008), random amplified polymorphic DNA (RAPD) (Ruwali et al., 2013), simple sequence repeat (SSR) (Montilla-Bascón et al., 2013), and single nucleotide polymorphisms (SNP) (Esvelt Klos et al., 2016). While these approaches have advanced our understanding of oats genetics globally, there remains a significant gap in comprehensive genetic diversity for oat in Ethiopia. In particular, the application of SNP markers, now regarded as the gold standard for high-resolution diversity and population analysis in crop species has been limited in Ethiopian germplasm.

While assessing genetic diversity is foundational for conservation and pre-breeding, understanding the genetic architecture of key traits is equally critical for developing improved oat varieties. Most economically important agronomic traits such as yield, biomass and stress tolerance are quantitatively inherited and controlled by numerous small-effect genes or quantitative trait loci (QTLs), with their expression significantly influenced by genotype × environment interactions (Bhat et al., 2021; Boopathi et al., 2022; Farooqi et al., 2022). Dissecting the genetic basis of complex traits is essential for accelerating breeding efforts aimed at enhancing adaptation to diverse and changing climatic conditions (Khan et al., 2021). Genome-wide association study (GWAS) has emerged as a powerful approach for unravelling the genetic architecture of these complex traits, enabling the identification of genomic regions associated with phenotypic variation at relatively high resolution (Wang et al., 2020; Uffelmann et al., 2021). In oats, GWAS has been successfully applied to uncover loci associated with traits such as β-glucan content (Zimmer et al., 2020; Bazzer et al., 2025), lemma color (Winkler et al., 2016; Wang et al., 2023), and disease resistances, particularly to crown rust (Klos et al., 2017; Hewitt et al., 2024). However, despite the growing interest in oat forage use, key agro-morphological traits relevant to forage performance, such as leaf length, tiller number, plant height, total fresh weight, and stem thickness, remain understudied. Only a limited number of GWAS studies have addressed these traits (Li et al., 2025; Peng et al., 2025) and many were conducted before the release of a fully annotated reference genome, restricting both their mapping precision and functional interpretability.

To address these gaps, the present study was designed with the following objectives: (1) to assess the genetic diversity of oat accessions, mainly from Ethiopia, using SNP markers; (2) to identify and map genomic regions significantly associated with key agro-morphological traits relevant to the dual-usage of forage and grain production; and (3) to identify high-yielding and phenotypically stable genotypes across the lesser studied subtropical environments, with the ultimate goal of supporting locally-targeted breeding programs focused on improving yield and feed quality traits in oats for the needs of the East African stakeholders.

## MATERIAL AND METHODS

### Plant materials

This study utilized a panel of 169 oat (*Avena sativa*) accessions, comprising 119 obtained from the Ethiopian Agricultural Research Institute (EIAR), Holetta Agricultural Research Center (HARC), and 50 from the International Livestock Research Institute (ILRI) genebank with more broader origins as standard checks or controls (Supp. table 1).

### Field Experiment: Experimental Design and Site Description

The field experiment was conducted over two consecutive main cropping seasons (2022 and 2023) across three different locations in Ethiopia. The first site, Holetta, is located at 9°03’ N, 38°30’ E, at an altitude of 2390 meters above sea level (m.a.s.l.), and receives an average annual rainfall of 1091.51mm. The soil type is Nitosol soil, and the site experiences maximum and minimum temperatures of 22.2°C and 6.13°C, respectively. The second site, Adda Berga, lies at 9°18’ N, 38°28’ E, at an altitude of 2592 m.a.s.l. and receives an average annual rainfall of 1225 mm. The maximum and minimum temperatures recorded at this location are 25°C and 10°C, respectively. The third site, Addis Ababa, is situated at 8°58’ N, 38°45’ E, at an elevation of 2450 m.a.s.l., with annual rainfall of 1089 mm and temperatures ranging from 22°C to 8.3°C.

The field experiment was laid out in a partially balanced simple lattice design (13*13) with two replications at each location. Seeds were directly sown in 5.2 meters-long rows, spaced 20 cm apart. Each incomplete block consisted of 26 rows, with each oat accession represented by two rows. Spacing between incomplete blocks and replications was maintained at 1 meter and 1.5 meters, respectively. A seeding rate of 100 kg per hectare was used, and NPS fertilizer (a mix of Nitrogen, Phosphorus, and Sulfur) applied at a rate of 100 kg per hectare at sowing. Weed management was carried out manually through hand weeding, starting from plant emergence and repeated at two-week intervals until harvest.

### Phenotypic Data Collection

Plant vigor (Vig_score) was visually assessed two weeks after seedling emergence using a standardized scoring scale to capture early growth performance. Four morphological traits, plant height (PH), leaf length (LL), stem thickness (ST), and number of leaves per plant (NLPP) were evaluated at the soft dough stage (Tottman, 1987). For each accession, measurements were taken from five randomly selected plants and their average was recorded. PH and LL were measured using a measuring tape, ST with a caliper, while NLPP was determined by counting the number of leaves per plant (supp table 2). For biomass-related traits, total fresh weight (TFW, kg) was recorded immediately after harvesting at the soft dough stage while oven dry weight (ODWT, g) was determined by drying a subsample at 60 °C for 72 h to a constant weight. Fresh yield per hectare (FTha) was estimated by converting the total fresh weight (TFW) from the harvested plot area using the formula and dry matter yield per hectare (DTha) was then estimated based on green forage yield and the dry matter concentration of the oven-dried samples. The dried subsamples were subsequently used to assess forage quality traits. Grain quality traits were evaluated using seeds harvested at maturity stage, dried, and threshed. Traits including seed area (SA), seed width (SW), seed length (SL), and thousand-grain weight (TGW) were measured on hulled seeds using a Marvin seed analyzer (MARViTECH GmbH, Germany) (Supp table 2).

#### Feed Quality Analysis

Oven-dried tissue samples from whole plants, leaves, and stems were ground separately to pass through a 1 mm sieve. The finely ground samples were scanned using Near-Infrared Spectrometer (NIRS) (FOSS Forage Analyzer 5000; software package WinISI II). Previously developed oat equation was used to predict the spectra to estimate feed quality traits following AOAC procedure “Fiber (Acid Detergent) and Protein (Crude) in Animal Feed and Forages: Near-infrared Reflectance Spectroscopic Method. (989.03) Official Methods of Analysis. 1990. Association of Official Analytical Chemists. 15th Edition.”. The traits assessed included acid detergent fiber (ADF), neutral detergent fiber (NDF), acid detergent lignin (ADL), organic matter (OM), dry matter (DM), inorganic matter (ash), total nitrogen (N), crude protein (CP), in vitro organic matter digestibility (IVOMD), and metabolizable energy (ME). The predicted values were adjusted for DM, and percentage values were used for statistical analyses. Metabolizable energy yield (MEY) was calculated as the product of ME and total dry weight (TDW).

#### Genomic DNA extraction, GBS Library Preparation, and Sequencing

Genomic DNA was extracted from freeze-dried young leaf samples (stored at a -80 °C for 48 hours) using a DNeasy Plant Small Kit (Qiagen Inc., Valencia, CA), following the manufacturer’s protocol. DNA quantity and concentration were assessed using a Thermo Scientific NanoDrop Spectrophotometer (DeNovix DS-11 FX), and DNA integrity was verified by electrophoresis on a 1% agarose gel run in 1% TAE buffer at 100 V for 45 minutes.

For GBS library preparation, the restriction enzymes Pstl and ApeKl were selected based on recommendations from LGC Biosearch Technologies and their combination produced a fragment size suitable for Illumina sequencing. Paired-end sequencing (2×150 bp) was performed on an Illumina NextSeq 500/550 v2 system using Pstl-ApeKI site-specific primers. Sequencing libraries from each lane was demultiplexed using the samples’ barcode information for sample identification and Illumina’s bcl2fastq v2.20 software. Adapter sequences were trimmed during processing. Reads containing ambiguous nucleotides (Ns), the expected restriction sites, or shorter than 20 bases were discarded. Quality control was performed using FastQC (Andrews, 2010) and read count summaries were compiled.

#### SNP Calling and Data Filtering

Adaptor clipped reads were mapped to the *A. sativa* cv. Sanfensan reference genome (Peng et al., 2022) using the Burrows-Wheeler Aligner (BWA v0.7.17) (Li & Durbin, 2009). The resulting SAM files were converted into BAM format using SAMtools tools v1.15.1 (Li *et al.,* 2009) for downstream processing. Variant discovery and SNP calling were conducted using Next Generation Sequencing Experience Platform (NGSEP) version 4.2.1 (Tello et al., 2019). Initial SNP filtering retained SNP markers with a call rate of ≥90 % and minor allele frequency (MAF) ≥ 5% for downstream analysis. For genetic diversity analyses, markers were further refined based on Polymorphic information content (PIC) ≥ 0.2 and expected heterozygosity (He) ≥ 0.2. Missing genotype data were imputed using the Linkage Disequilibrium-based k-nearest neighbors (LD KNNi) algorithm implemented in TASSEL v5 (Bradbury *et al*., 2007), which uses LD patterns to identify the most similar SNPs and predict missing values accurately.

#### Phenotypic data analysis

To evaluate trait variability among genotypes across the three locations, descriptive statistics, including mean, standard deviation (SD), minimum, maximum, and coefficient of variation (CV) were computed using R software version 4.3.1 (R Core Team, 2013). Additionally, genetic variability parameters, such as broad-sense heritability (H²), genetic advance (GA), and genetic advance as a percentage of the mean (GAM) were estimated using the following linear model:

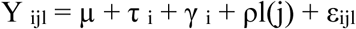

Where τ_i_ represents the treatment effect, γ_i_ denotes replicate effect, ρl(j) = block within replicate effect, and ε_ijl_=random error. Best linear unbiased predictions (BLUPs) for each trait were estimated using the mixed linear model in the “lme4” R package (Bates et al., 2015).

The combined analysis of variance (ANOVA) was also performed using the R package lme4, treating genotype as a fixed effect, and year, location, and block as random effects, according to the following model structure:

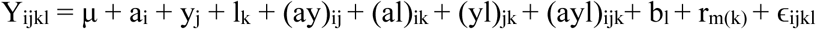

Where:

Y_ijkl_ is the observed response of genotype, μ is the overall mean, a_i_ is the effect of the i-th genotype, Y_j_ is the effect of the j-th year, l_k_ is the effect of the k-th location, bl is the effect of block, rm(k) is the replication effect nested within location, and ɛijkl is the residual error.

#### Principal Component and Cluster Analysis from phenotypes

Principal component analysis (PCA) and cluster analysis were conducted using R software version 4.3.1 software (R Core Team, 2013) to assess population structure and genetic relationships among the oat accessions. The optimal numbers of clusters and accession memberships were determined using the FactoMineR and factoextra R packages (Kassambara and Mundt, 2017). Clustering patterns were visualized through PCA biplots and dendrograms, providing insights into the genetic differentiation and groupings within the panel.

#### Genetic Diversity, Population Structure, and Phylogenetic Relationship analysis

Genetic diversity parameters, including expected heterozygosity (He) and polymorphic information content (PIC), were calculated for each SNP locus using the snpReady package in R software (R Core Team, 2013; Granato et al., 2018). Phylogenetic relationships among the oat accessions were assessed through hierarchical clustering based on a Euclidean distance matrix using MEGA version 6.0 (Tamura et al., 2013), with 1,000 bootstrap replicates to evaluate node support. Discriminant Analysis of Principal Components (DAPC) was performed using the adegenet package in R software (Jombart et al., 2018) to further explore population structure. The optimal number of clusters (K) was determined by identifying the minimum Bayesian Information Criterion (BIC) across a range of K values (1 to 40). Population genetic structure was also evaluated using Analysis of Molecular Variance (AMOVA) to partition genetic variation among and within clusters. AMOVA was conducted using the poppr package in R (Kamvar *et al*., 2014), providing insights into the distribution of genetic variation across the population.

#### Population Structure and Principal Component Analysis (PCA) from markers

Population structure was assessed using a Bayesian model-based clustering algorithm implemented in the STRUCTURE software version 2.3.4 (Pritchard, 2000), based on a subset of 1,823 robust SNP markers. To determine the optimal number of genetic clusters (*K*), a simulation was run with a burn-in period of 10,000 and 50,000 Markov Chain Monte Carlo (MCMC) iterations, for *K* values ranging from 1 to 10, with 10 independent replicates for each K. The most optimal number of genetic clusters (*K*) was inferred using ΔK method described by Evanno et al. (2005), as implemented in the web-based Structure Selector tool (Li and Liu, 2018). Accessions with a membership coefficient (Q-values) greater than 0.7 were assigned to a specific cluster, whereas those with Q-values below 0.7 were classified as admixed.

To complement STRUCTURE analysis, Principal Component Analysis (PCA) was performed to examine the genetic structure of the oat accessions. PCA was conducted in R software version 4.3.1 (R Core Team, 2013) using the FactoMineR package for calculate the PCA and factoextra for visualization of the results (Kassambara and Mundt, 2017).

#### Linkage Disequilibrium and Genome-wide Association Analysis (GWAS)

Pairwise linkage disequilibrium LD estimates, including the squared allele frequency correlations (r^2^) and associated *p*-values, were calculated between markers on each chromosome using TASSEL version 5.0 (Bradbury et al., 2007), under default parameters. The extent of LD decay across the genome was visualized by plotting the r^2^ values against physical distance (in base pairs) using R software (R Core Team, 2013). An r^2^ threshold of 0.2 was used as the baseline to define the point of linkage equilibrium between marker pairs.

Genome-wide association study (GWAS) was performed to identify significant associations between phenotypic traits from 167 oat genotypes. Two accessions with more than 50% missing genotype data were excluded, resulting in 167 genotypes used for this analysis. A total of 18,196 high-quality biallelic SNP markers (MAF > 5 %, missing values < 10%) were retained for the analysis. Significantly associated loci were identified based on a false discovery rate (FDR) adjusted *p*-value threshold of < 0.05.

A marker-trait association analyses were conducted using both single-locus and multi-locus models including: General Linear Model (GLM), Mixed Linear Model (MLM) (Zhang et al., 2010), Multi-Locus Mixed Model (MLMM) (Segura et al., 2012), Fixed and random model Circulating Probability Unification (FarmCPU) (Liu et al., 2016), and Bayesian-information and Linkage-disequilibrium Iteratively Nested Keyway (BLINK) (Huang et al 2019). These models were implemented in the Genome Association and Prediction Integrated Tool (GAPIT) in R (Tang et al., 2016).

The GWAS results were visualized using Manhattan plots in R with the ggplot2 package (Wickham et al., 2016), where the -log10 (*p*) values of SNPs were plotted against its genomic positions to identify significant association peaks.

#### Candidate Gene Identification and Functional Annotation

To identify potential candidate genes associated with the significant SNP-trait associations, the *Avena sativa* cv. Sanfensan reference genome was used. Genes located within a 200-kilobase (kb) window upstream and downstream of each significant SNP were considered putative candidates. Information on gene proximity and annotation details was retrieved using the Ensemble Plants genome browser. Functional characterization of these candidate genes was further conducted using the Oat Bioinformatics Database (OatBioDB: (http://www.waooat.cn), to explore gene functions, biological pathways, and relevance to agronomic traits.

## Results

### Phenotypic Variation

Most traits displayed an approximately normal distribution across the three tested environments. Notably, seed-related traits such as seed width (SW), seed length (SL), seed area (SA) and forage traits like stem thickness (ST), and number of leaves per plant (NLPP), and leaf length (LL) exhibited a consistent normal distribution across all locations (Supplementary Figure 1). Descriptive statistical analysis revealed environmental effects on traits like total fresh weight (TFW), fresh weight per hectare (FTha), dry weight per hectare (DTha), and plant height (PH), with the highest mean values for these traits predominantly recorded at the Holetta site. Broad-sense heritability (H²) estimates ranged from low (0.09 for oven dry matter (DM)) to high (0.77 for seed width (SW)), indicating varying degrees of genetic control across traits. The genetic advance as a percentage of mean (GAM) also varied, ranging from 0.09% to 34.74%, with seed-related traits exhibiting the highest genetic gains (Table 1), suggesting strong potential for selection and improvement in these traits. In contrast, feed quality traits exhibited the lowest values for both H² and GAM, suggesting limited prospects for genetic improvement in these traits.

**Table 1:**
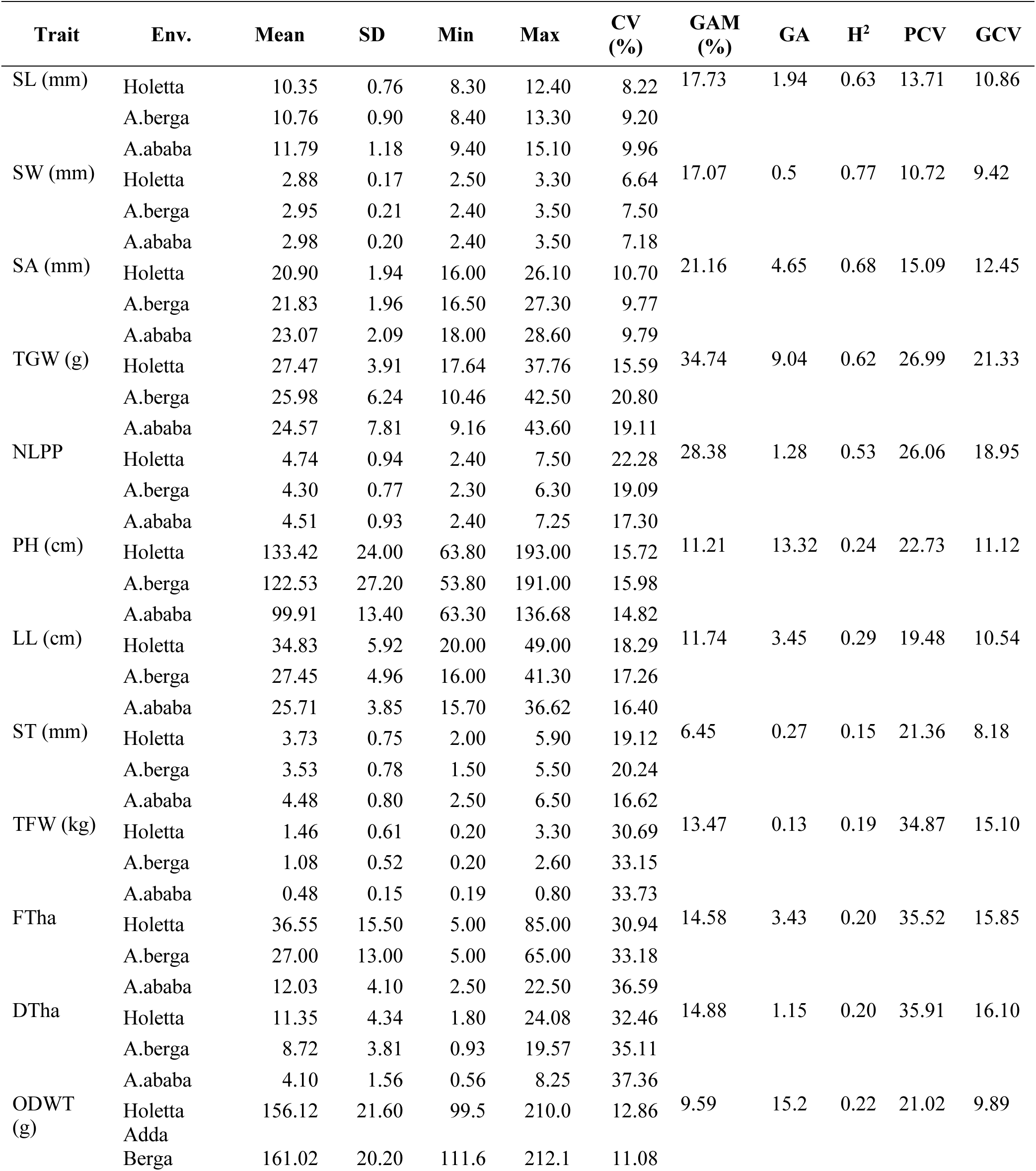
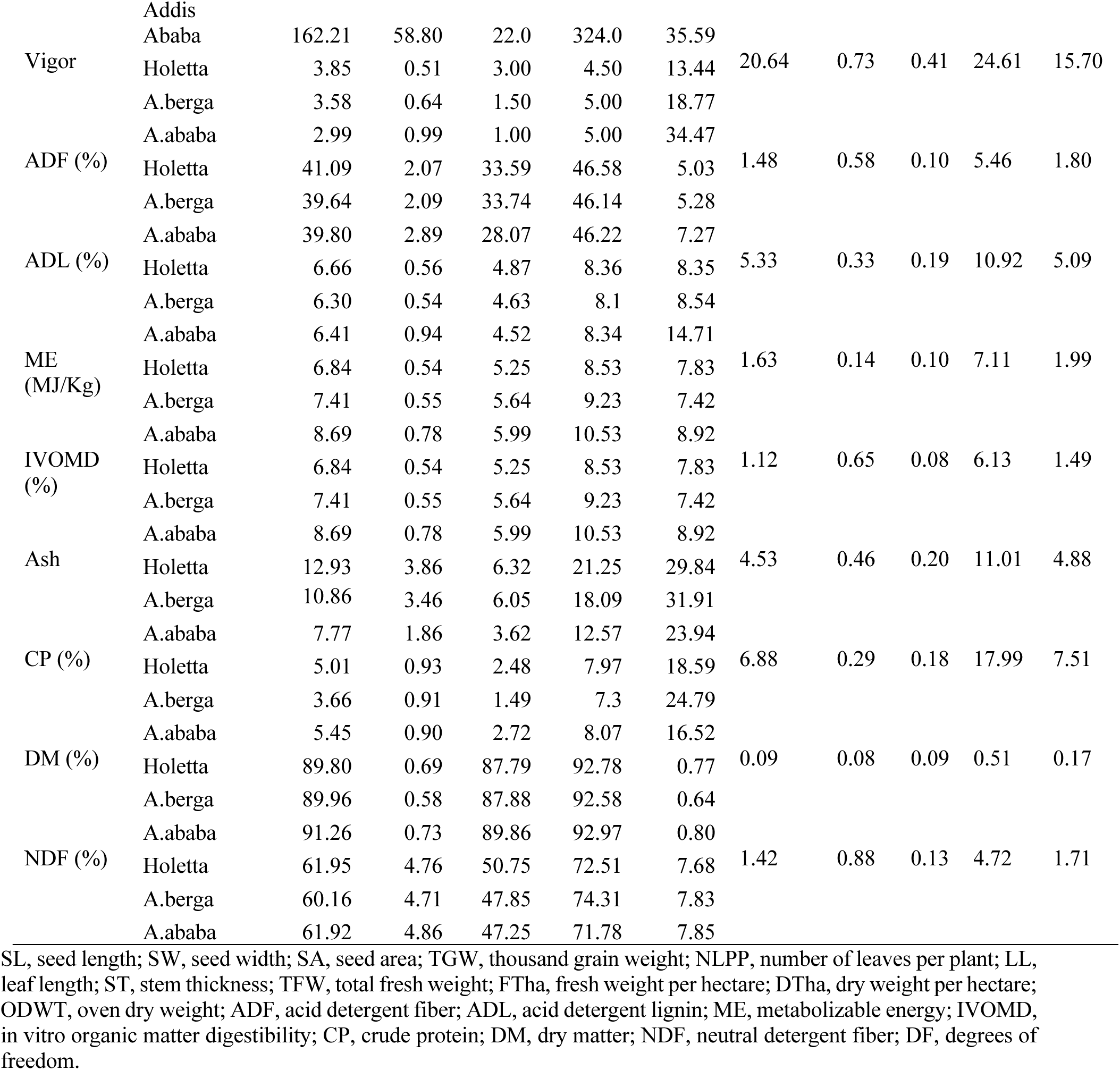
Descriptive statistics, broad-sense heritability (H^2^), genetic advance (GA), and genetic advance as percentage of mean (GAM), genotypic coefficient of variance (GCV), and phenotypic coefficient of variance (PCV) for biomass, feed quality, and grain-related traits evaluated across three environments.

**Table 2:**
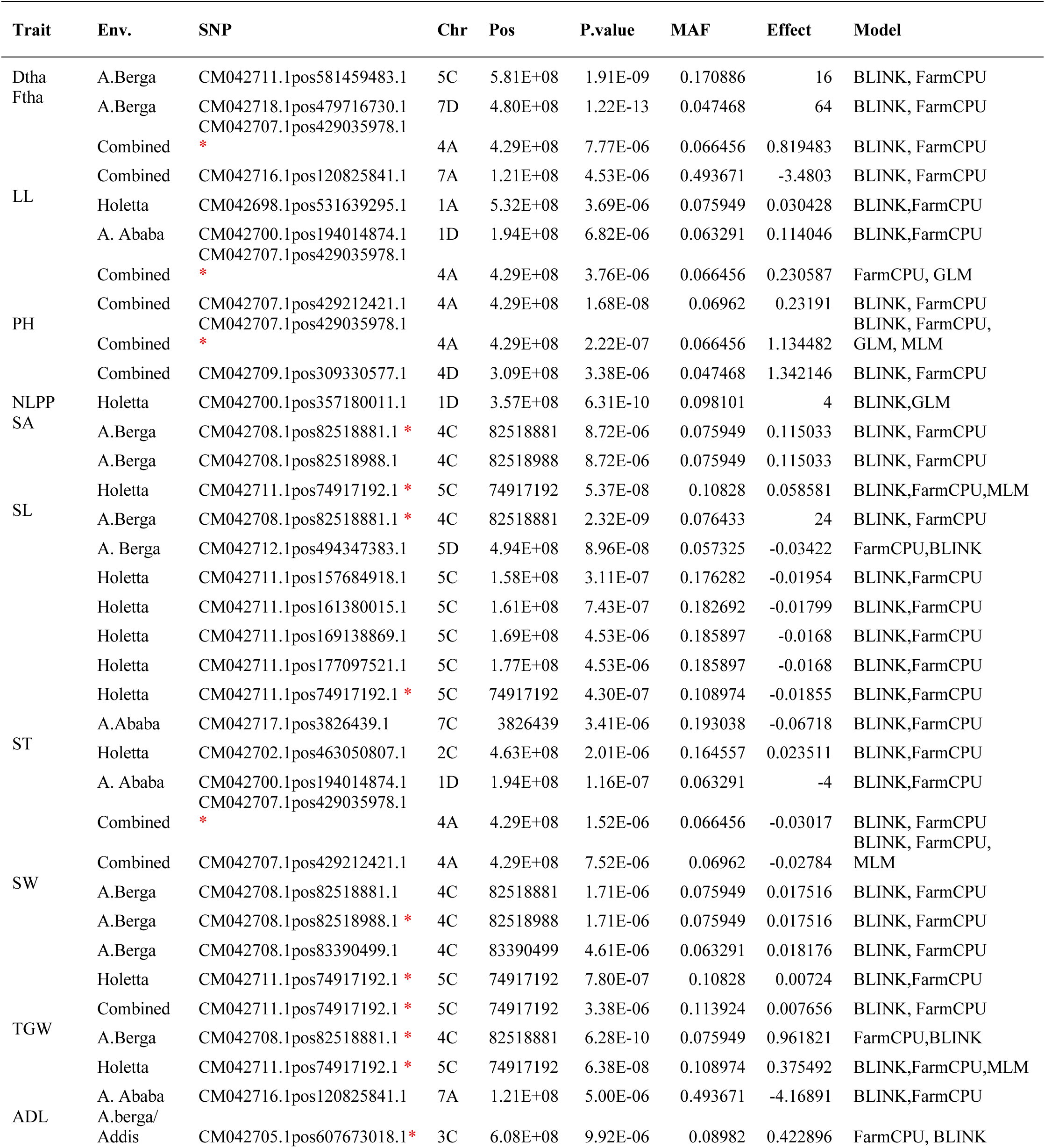
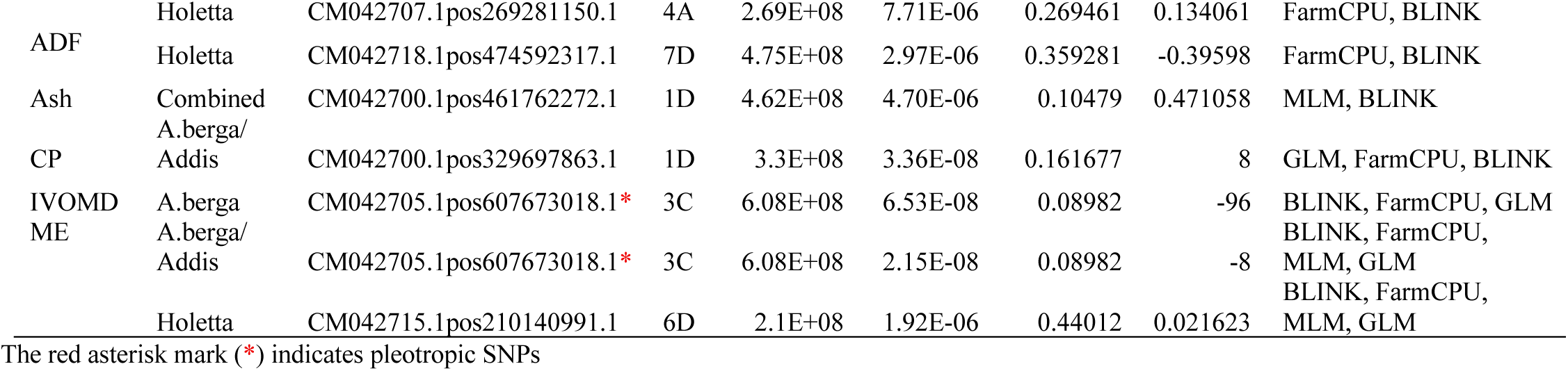
Summary of genome-wide analysis (GWAS) results for selected agronomic traits. The table includes information on chromosome location, physical position, SNP ID, minor allele frequency (MAF), allelic effect, and the GWAS models in which the association was detected.

The phenotypic coefficient of variation (PCV) was consistently higher than the genotypic coefficient of variation (GCV) for all traits, indicating a substantial environmental influence on trait expression. However, relatively comparable GCV values were observed for grain-related traits, particularly SW, SA, and SL (Table 1).

The combined ANOVA across the three locations revealed highly significant effects of genotype and year (p < 0.0001) for all evaluated traits, highlighting the strong influence of genetic and temporal factors on trait expression. Environmental factors were also significant for most traits, with the exception of ODWT, suggesting differential trait responsiveness to location-specific conditions. Significant two-way interaction effects (genotype × year, genotype × environment, and year × environment) were also observed for most of traits, excluding vigor score, SW and LL, indicating variable stability of these traits across conditions. Additionally, the three-way interactions (genotype × year × environment) were also significant (p < 0.05 or p < 0.001) for most traits, underscoring the complexity of genotype-by-environment interactions and the need for multi-environment testing to identify broadly adapted genotypes (Supp. table 3).

### Principal Component and Cluster Analysis based on Phenotypic Traits

To explore the underlying structure of phenotypic variation among oat accessions multivariate analyses, including Principal Component Analysis (PCA) and hierarchical cluster analysis were conducted. PCA was used to reduce data dimensionality by identifying combination of traits that contributed most to the overall variation, while cluster analysis grouped genotypes based on their overall similarity. The first three principal components (PCs) accounted for a cumulative 51.1% of the total phenotypic variance (Supp. table 4). PC1 explained the largest proportion (24.28%) and was primarily influenced by feed quality traits, including ADL (0.42), ADF (0.41), and NDF (0.39). PC2 explained 17.02% of the variation and was associated predominantly with seed-related traits such as TGW, SA, and SW. PC3 contributed 10.42% of the variation, with high loadings for ST (0.38) and ODWT (0.32). The PCA biplot (Figure 1A) illustrates correlations among traits, where the angle between vectors indicated the strength and direction of association. Feed quality traits, ADF, NDF, and ADL, cluster together with vectors projecting towards the positive Dim1 axis and exhibit strong positive correlations among themselves. In contrast, IVOMD and ME are oriented in the negative Dim1 direction and display negative correlations with the fiber traits (ADF, ADL, and NDF). Grain-related traits TGW, SW, and SA form a distinct cluster upwards, showing positive intercorrelations and clear separation from feed quality attributes, while also presenting negative correlations with biomass traits such as PH, ODWT, and Ftha. Other agronomic traits, including vigor, LL, and NLPP, contributed minimally to variation along the Dim1 and Dim2 axes (Figure 1A).

**Figure 1:**
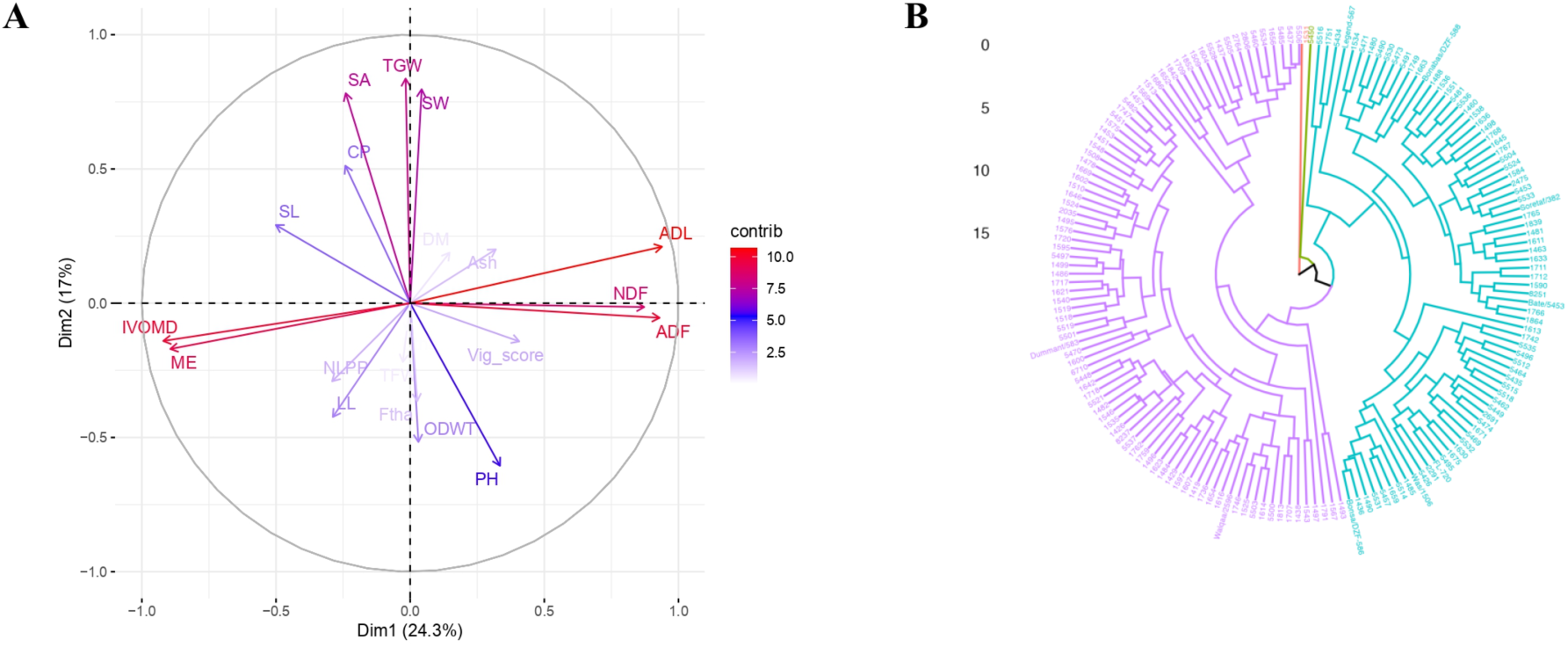
A) Principal component analysis plot showing the relationships among twenty-one traits in 167 oat accessions. B) Cluster Analysis of 167 oat accessions based on phenotype data.

Hierarchical cluster analysis grouped the oat accessions into two major clusters based on the mean values of twenty-one phenotypic traits (feed quality, grain-related, and agronomic traits), assessed across two years (Figure 1B). Notably, two genotypes (1531 and 5450) formed distinct singleton clusters, thereby highlighting their phenotypic divergence from all other genotypes.

### SNP Diversity and Chromosomal Distribution in oat Accessions

A total of 62,889 high-quality SNP markers were identified across 167 samples of oat (*Avena sativa*) accessions using Illumina NextSeq 500 platform. Of these, 62,221 (98%) were successfully mapped to the 21 chromosomes of the *A.sativa* “Sanfensan” reference genome (Peng *et al.,* 2022), averaging 2,963 SNPs per chromosome (Figure 2A). Chromosome 4D had the highest number of SNPs (4,174), while chromosome 6D had the fewest (1,575). Among the subgenomes, the D genome exhibited the highest marker density (21,867), followed by the A and C genomes, with 20,952 and 19,402 markers, respectively (Figure 2A). An additional 668 SNPs (1.2%) were mapped to unanchored contigs.

**Figure 2:**
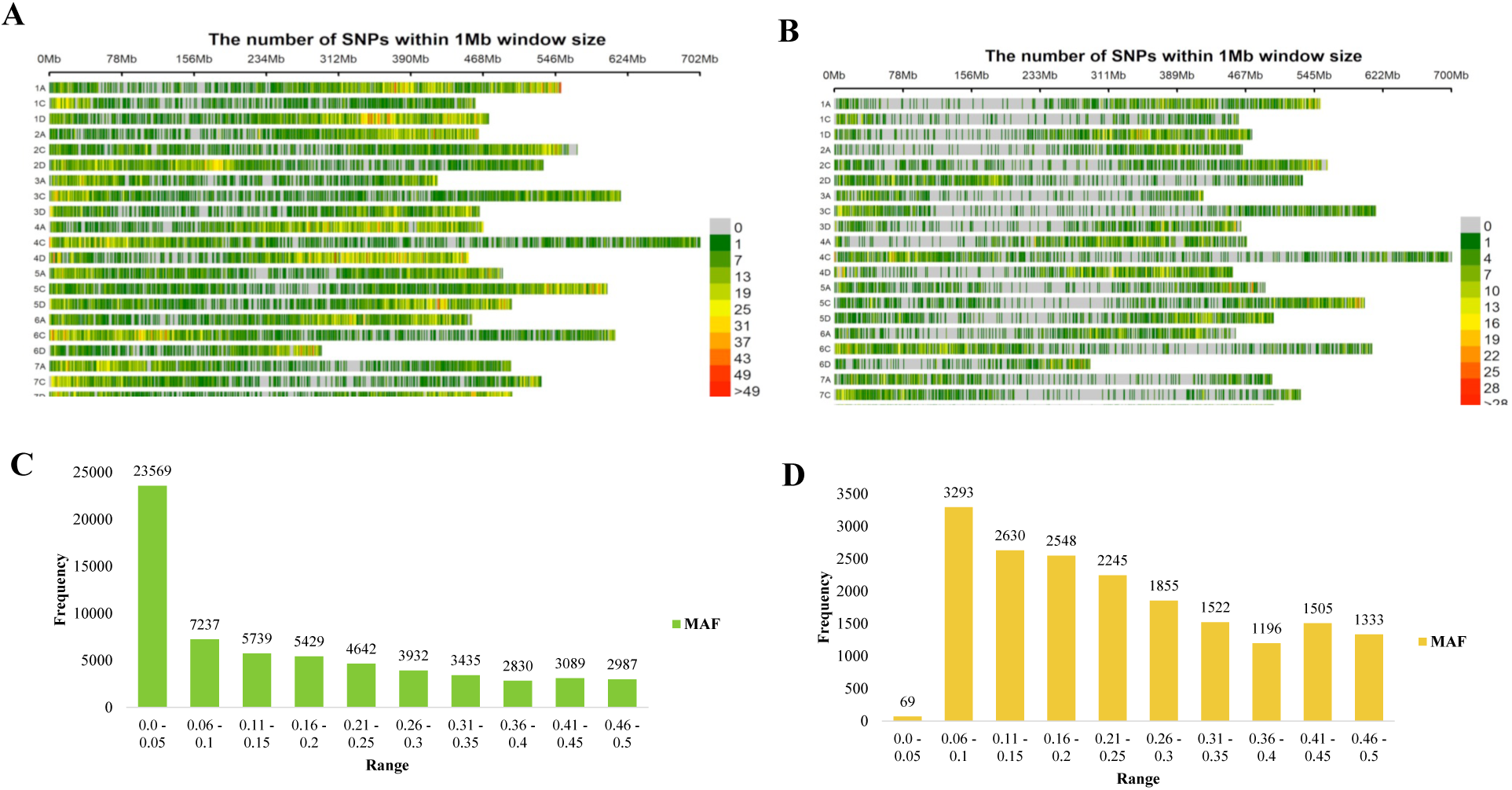
Density and distribution of single nucleotide polymorphisms (SNPs) across oat chromosomes, using 1 Mb window size, and distribution of minor allele frequency (MAF). (A) SNPs density and chromosomal distribution before filtering. (B) SNP density and distribution after filtering. (C) MAF distribution prior to filtering. (D) MAF distribution after filtering.

The expected heterozygosity (He) of the SNP markers ranged from 0 to 0.5, with an average of 0.39 and a median of 0.46. The polymorphic information content (PIC) value varied from 0 to 0.38, with a mean of 0.3. Notably, approximately 69.7% of the genetic markers showed PIC values in the range of 0.3 to 0.38 (Supplementary figure 2A), indicating a high level of informativeness for most SNPs. The minor allele frequency (MAF) of SNPs ranged from 0.01 to 0.5, with an average of 0.15. Overall, 62.5% of the SNPs had MAF values greater than 0.05, suggesting adequate allelic diversity for reliable population structure and association analyses (Figure 2C). The proportion of markers’ missing data ranged from 0 to 40.8%, with a mean of 14.56%. Two accessions with more than 50% missing data were excluded, resulting in a final dataset of 167 accessions for downstream analysis.

After quality filtering, a total of 18,196 SNP markers with less than 10% missing data and a minor allele frequency (MAF) greater than 5% were retained for GWAS analysis. For population structure and genetic diversity analyses, a subset of 1,823 informative SNPs was selected based on thresholds of PIC ≥ 0.2 and He ≥ 0.2. These markers had an average He of 0.41 and a mean PIC value of 0.33 (Supplementary figure 2B). Of the 1,823 SNPs, 696 were mapped to the A genome, 331 to the C genome, and 796 to the D genome.

### Population Structure and Genetic Diversity Analysis

The maximum ΔK value supported *K*=2, as the most likely number of major clusters, while the peak at K=7 suggested the presence of seven subclusters (Supplementary figure 3). Out of 167 accessions, 135 (80.8 %) were confidently allocated to one of the two clusters (Pop1 and Pop2), with a membership probability threshold of > 70%, while the remaining 32 accessions (19.2 %) were classified as admixed. Pop1 represented the majority group, (71.3% of all accessions), and Pop2 comprised 9.6% (Figure 3A; Supp. table 5). Pop1 included a mix of EIAR (78 accessions, 65.5%) and ILRI (41 accessions, 34.5%) materials, while Pop2 was predominantly composed of EIAR accessions (14 of 16; 87.5%) with only two accessions from ILRI. Overall, the population structure analysis revealed the presence of two distinct genetic groups with moderate levels of admixture. Clustering patterns largely reflect the origin of the accessions, suggesting partial differentiation between EIAR and ILRI collections, and indicating some historical gene flow or shared ancestry among subsets of accessions.

**Figure 3:**
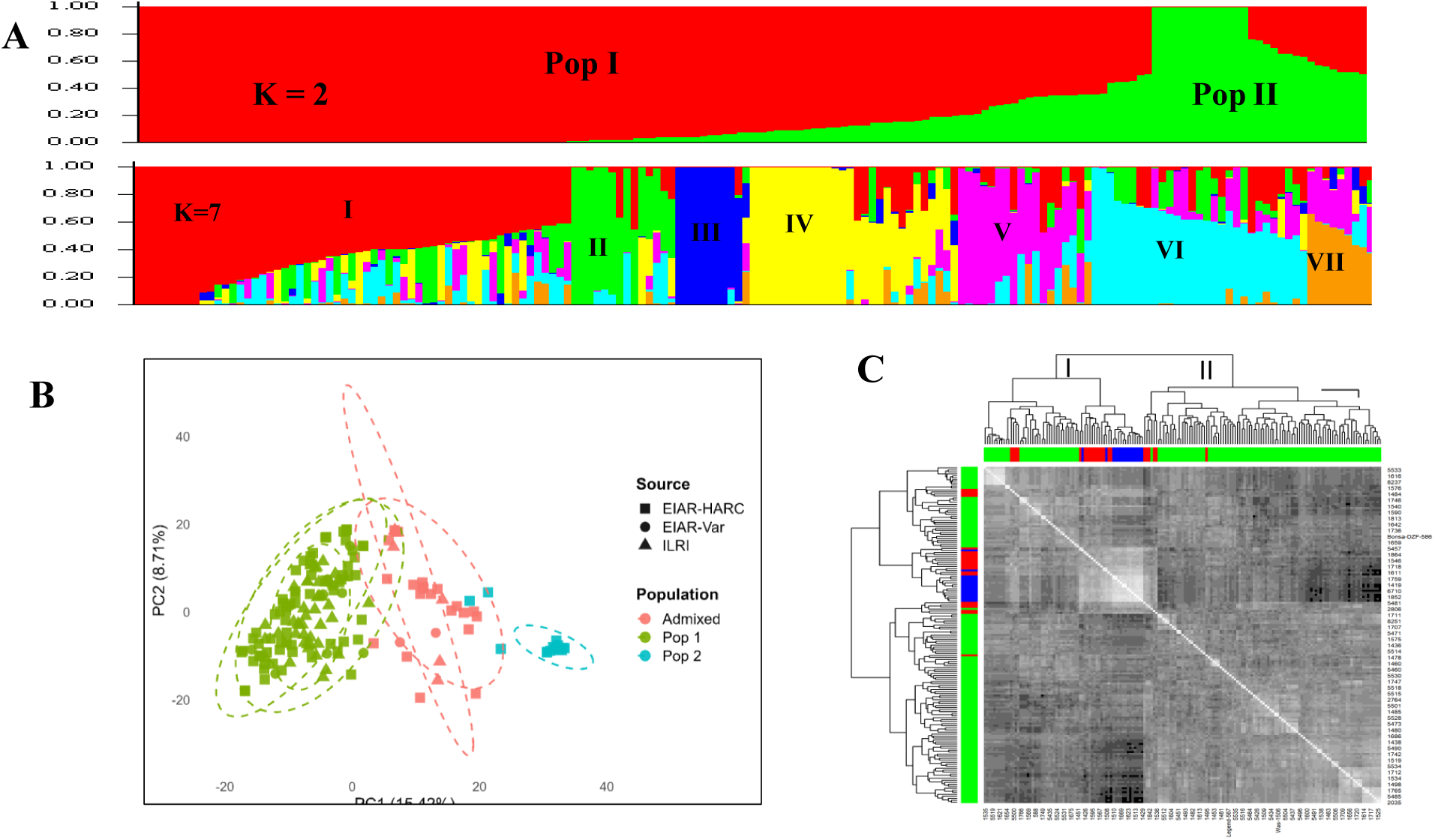
Population structure of 167 oat accessions based on 1823 SNP data. (A) Bar plots illustrating membership at *K* = 2 and at *K* = 7, with colors indicating assignments to subpopulations (B) Principal component analysis (PCA) plot, illustrating genetic subgroups identified by STRUCTURE. (C) A heat map of kinship matrix illustrating the genetic relationship between 167 oat accessions.

PCA further supported the genetic stratification observed in the STRUCTURE analysis. The first three principal components accounted for 29% of the total genetic variation with PC1 explaining 15.4%, PC2 8.7%, and PC3 4.9%. The PCA scatter plot clearly separated into two major cluster corresponding to pop1 and pop2, consistent with the results from the STRUCTURE analysis at K=2 (Figure 3A). This concordance between PCA and STRUCTURE highlights the robustness of the population differentiation and reinforces the presence of two primary genetic groups within this oat panel.

Kinship analysis further corroborated the population structure results, revealing two primary genetic clusters within this oat panel. These clusters were consistent with the STRUCTURE classification, with most accessions grouped within or near the same clusters identified by STRUCTURE. Furthermore, seven sub-clusters were identified, indicating finer scale genetic relationships among the accessions (Figure 3C). The heatmap, shaded by grayscale intensity, reflects pairwise kinship coefficients, with higher tones indicating genetic relatedness. Notably, several accessions, from Pop 2 exhibited strong genetic similarity, forming a distinct, closely related group within the boarder structure (Figure 3C).

The neighbor-joining (NJ) analysis classified the oat genotypes into three distinct clusters: C1, C2, and C3 (Figure 4A). Cluster C1 contained the largest number of accessions (71), followed by C2 with 62, and C3 with the smallest cluster, with 34 accessions (Supp. table 6). Most of the accessions assigned to Pop1 in the STRUCTURE analysis were grouped within C1, while all accessions from Pop2 clustered in C2. The remaining accessions Pop1 accessions, along with those classified as admixed in the STRUCTURE analysis, were predominantly found in C3. Consistent with the hierarchical clustering, Discriminant Analysis of Principal Components (DAPC) based on 1,823 SNPs, also grouped the 167 oat genotypes into three distinct clusters (Figure 4B). A scatter plot, using the three clusters and 200 principal components, was generated to visualize the spatial distribution of each genotype (Figure 4C).

**Figure 4:**
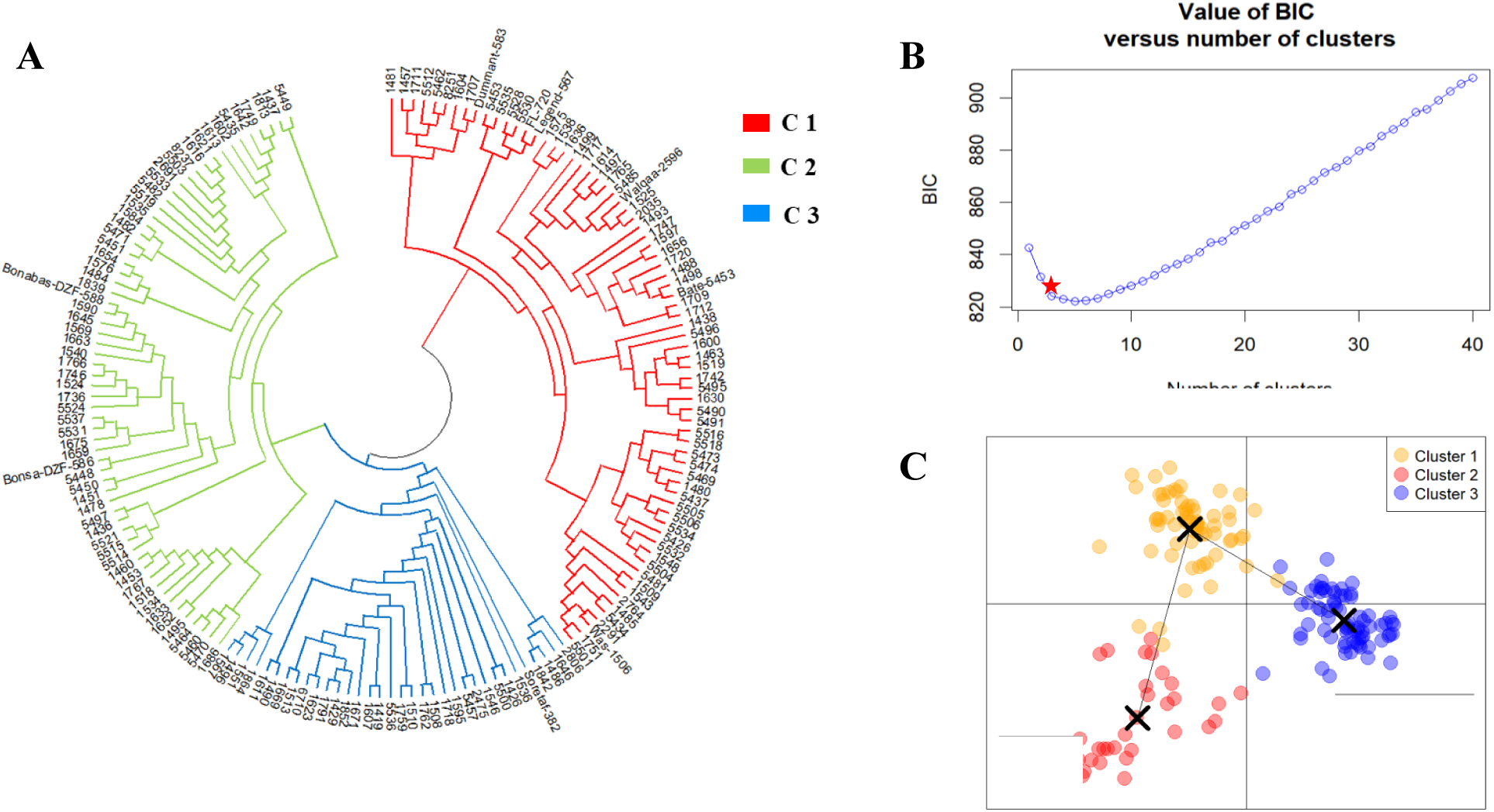
Hierarchical clustering and a discriminant analysis of principal components (DAPC) of 167 oat genotypes based on 1,823 SNP markers. (A) Dendrogram generated by hierarchical clustering using the neighbor-joining method with 1000 bootstrap replicates. Each genotype is represented by a branch, and the tree is divided into three colored subclades corresponding to the identified clusters. (B) Optimum number of clusters indicating K=3and (C) Scatterplot of the genotypes using the first two principal components distinguish the three clusters.C1= cluster 1, C2=cluster 2, and C3=cluster 3.

An analysis of molecular variance (AMOVA) was performed to assess genetic differentiation among groups defined by STRUCTURE-based subpopulations, hierarchical clustering, and growth habit classification. AMOVA based on population structure showed that 83% of the total genetic variation resided within populations, while 16.9% was attributed to differences among populations, (PhiPT = 0.17; Supp. table 7). AMOVA based on hierarchical clustering explained the highest proportion of genetic variation among groups, accounting for 18.2% of the total variance and a PhiPT value of 0.18, supporting the findings from the population structure analysis. In contrast, AMOVA based on growth habit explained only 3.4% of the genetic variation among groups, with 96.6% occurring within groups, indicating minimal differentiation based on growth habit (Supp. table 7).

### Linkage Disequilibrium (LD) Analysis

Linkage disequilibrium (LD) patterns varied both across chromosomes and subgenomes. A total of 597,327 marker pairs were identified, with an average LD value (r²) of 0.25. The C genome accounted for the highest number of marker pairs (259,072; 43.4%), followed by the D genome (165,025; 27.6%) (Supp. table 8), while the A genome contributed the fewest (155,230 or 26%). The weakest average LD value (r² = 0.18) was observed among marker pairs on chromosomes 1C, 2D, and 6D, indicating relatively low linkage. In contrast, the strongest LD (r² = 0.35) was detected between marker pairs on chromosome 3D and 5A, occurring at an average physical distance of 21,782 and 18,397 bps, respectively (Supp. table 8). Across all chromosomes, LD decay began at r^2^ = 0.46 and reached half of this initial value at a physical distance of 7,369,873 bp between marker pairs (Supplementary figure 4). For QTL mapping, significant SNPs located on the same chromosome were assigned to a single QTL region if the physical distance between them was less than the 7,369,873 bp decay threshold.

### Marker-Traits Association Analysis across Multiple Environments

GWAS analysis was performed using 18,196 high-quality SNP markers and utilized Best Linear Unbiased Prediction (BLUP) values derived from three environments. Analyses were conducted separately for each environment and on the combined dataset, using both single-locus and multi-locus models to enhance detection accuracy. Initially, 186 SNPs showed significant association with various traits (Supp. table 9). To enhance robustness, only SNPs identified by at least two models were retained, resulting in 42 high-confidence marker-trait associations (MTAs) across 11 traits. These included Dtha, Ftha, LA, LL, PH, ST, NLPP, SA, SL, SW, and TGW. The number of associated SNPs per trait ranged from 1 to 8, reflecting polygenic nature of these traits (Table 2).

Among the 42 SNPs identified, four exhibited pleiotropic effects, being significantly associated with multiple traits. One such marker, CM042707.1pos429035978.1, (chromosome 4A) was linked to multiple agronomic traits (Plant height, stem thickness, fresh weight per hectare, and leaf length). Another pleiotropic marker, CM042708.1pos82518881.1 (chromosome 4C), and CM042711.1pos74917192.1 (chromosome 5C) were associated with seed area, seed weight, seed length, and thousand-grain weight. Notably, CM042711.1pos74917192.1 demonstrated consistent association with seed weight across two different environments, highlighting its potential for marker-assisted selection. Another SNP, CM042705.1pos607673018.1 (chromosome 3C), was also associated with three feed quality traits: acid detergent lignin (ADL), metabolizable energy (ME), and in vitro organic matter digestibility (IVOMD) (Figure 5).

**Figure 5:**
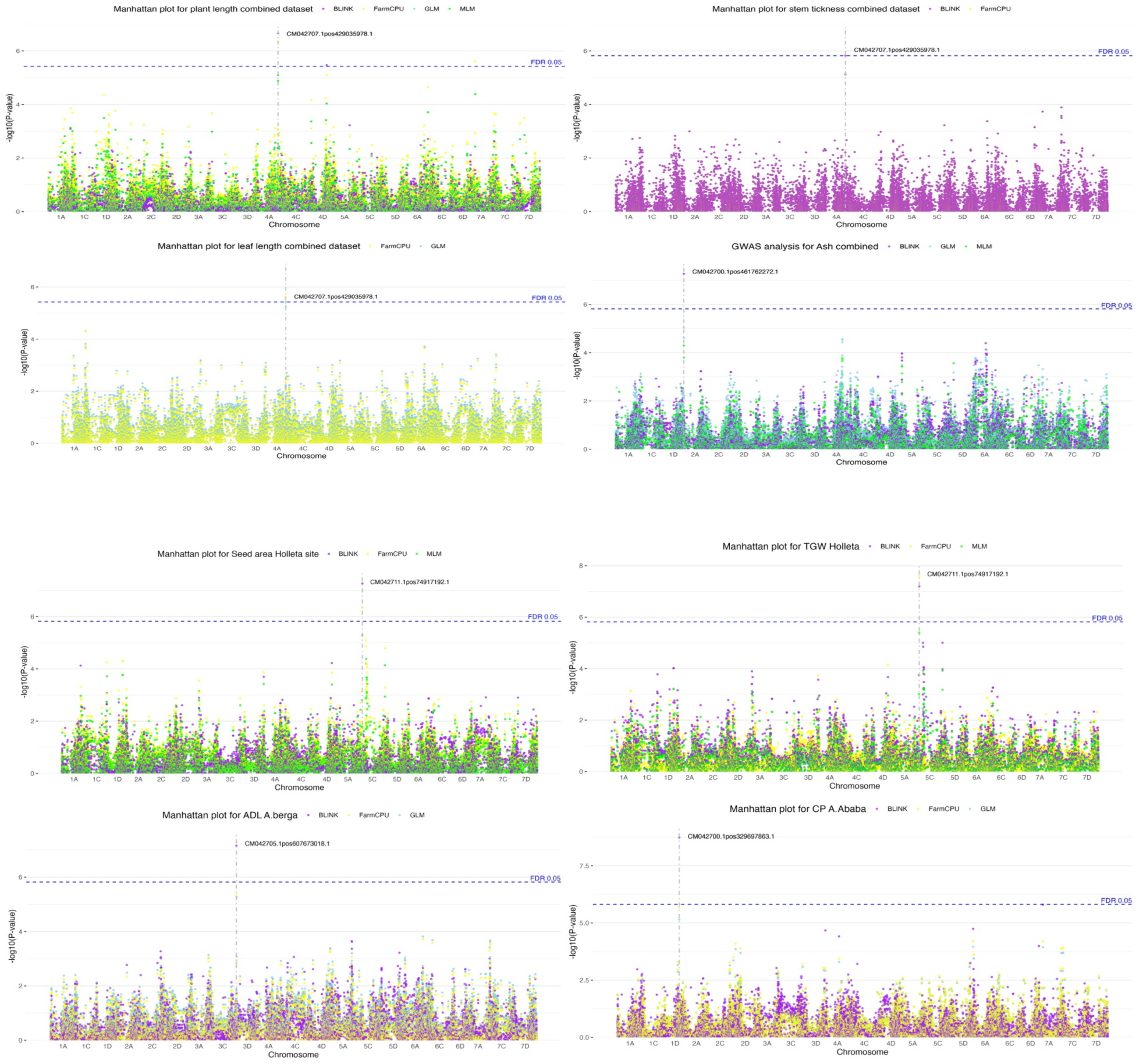
Manhattan plots illustrating GWAS results for selected traits. Each point represents a single nucleotide polymorphism (SNP), with the x-axis showing genomic positions across chromosomes and the y-axis indicating the significance of association with the trait.

### Candidate Gene Identification

A total of 46 candidate genes were identified within a 200 kb window flanking the associated SNP loci, with up to five genes located at each locus. These genes were annotated and classified according to their biological processes and molecular functions, and grouped into key functional categories, including enzymes, transcription factors, receptors and signal transduction proteins, transporters, and proteins involved in degradation and ubiquitination (Table 3). This functional diversity highlights the complex genetic architecture underlying the associated agronomic traits.

**Table 3:**
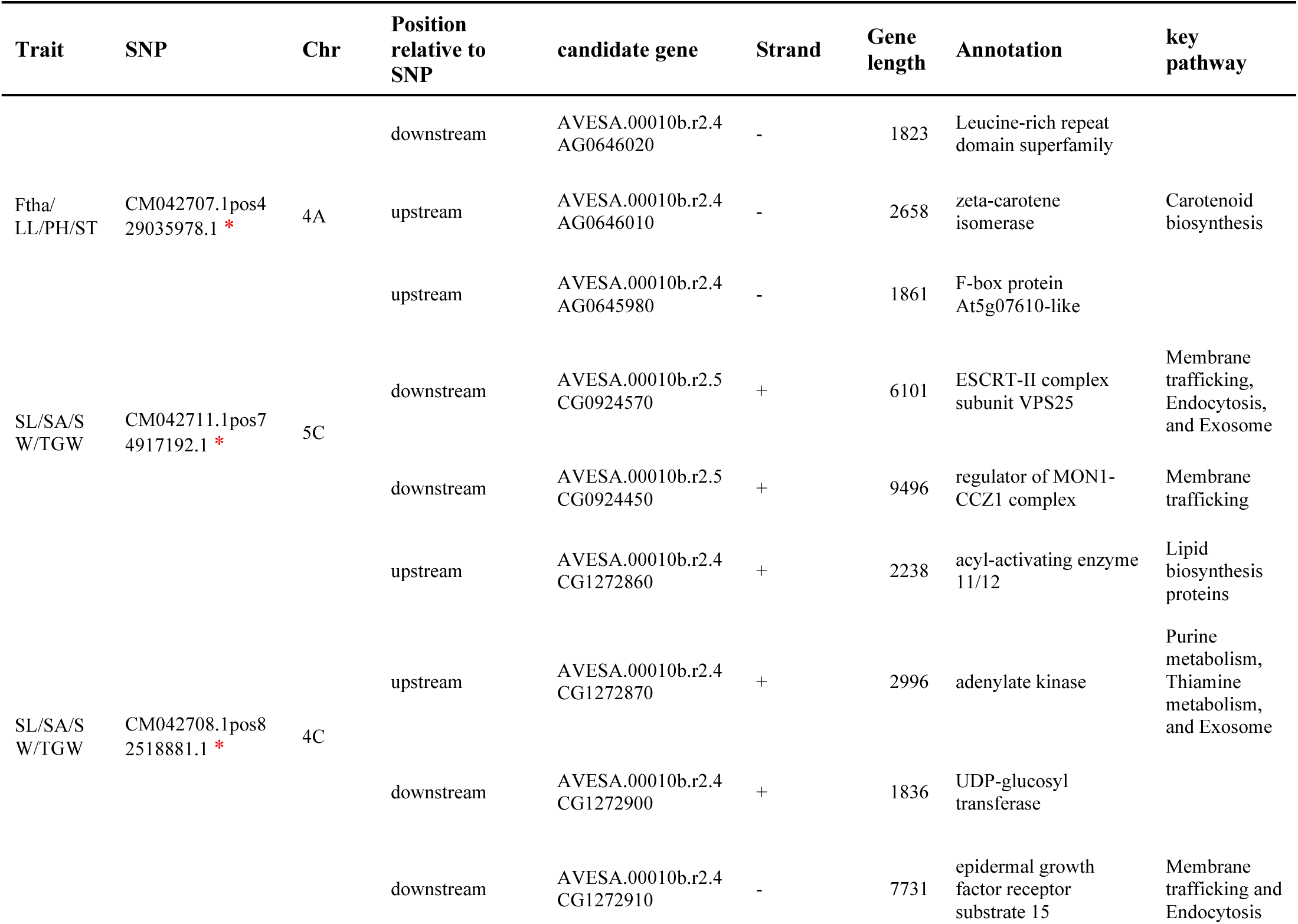
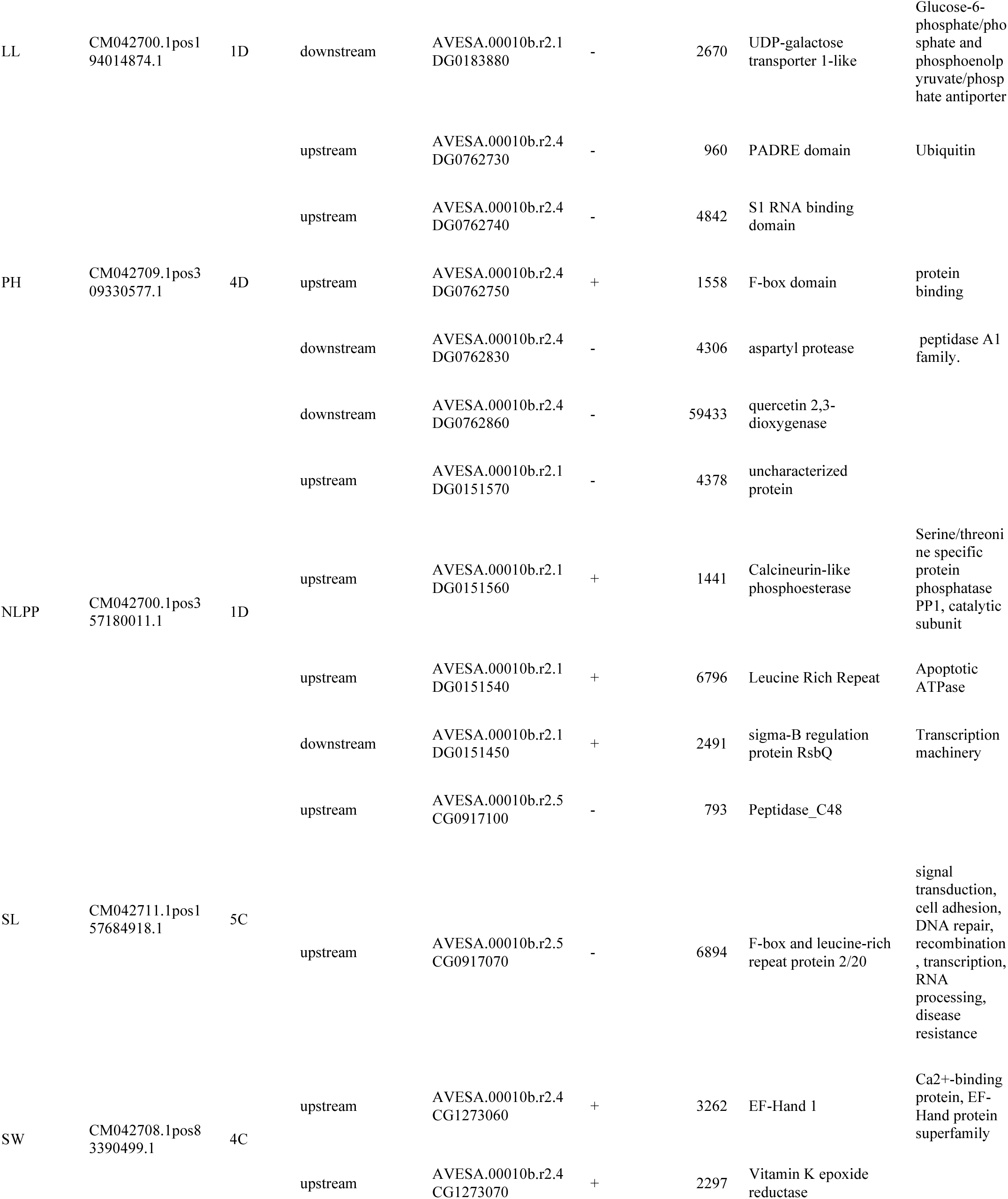
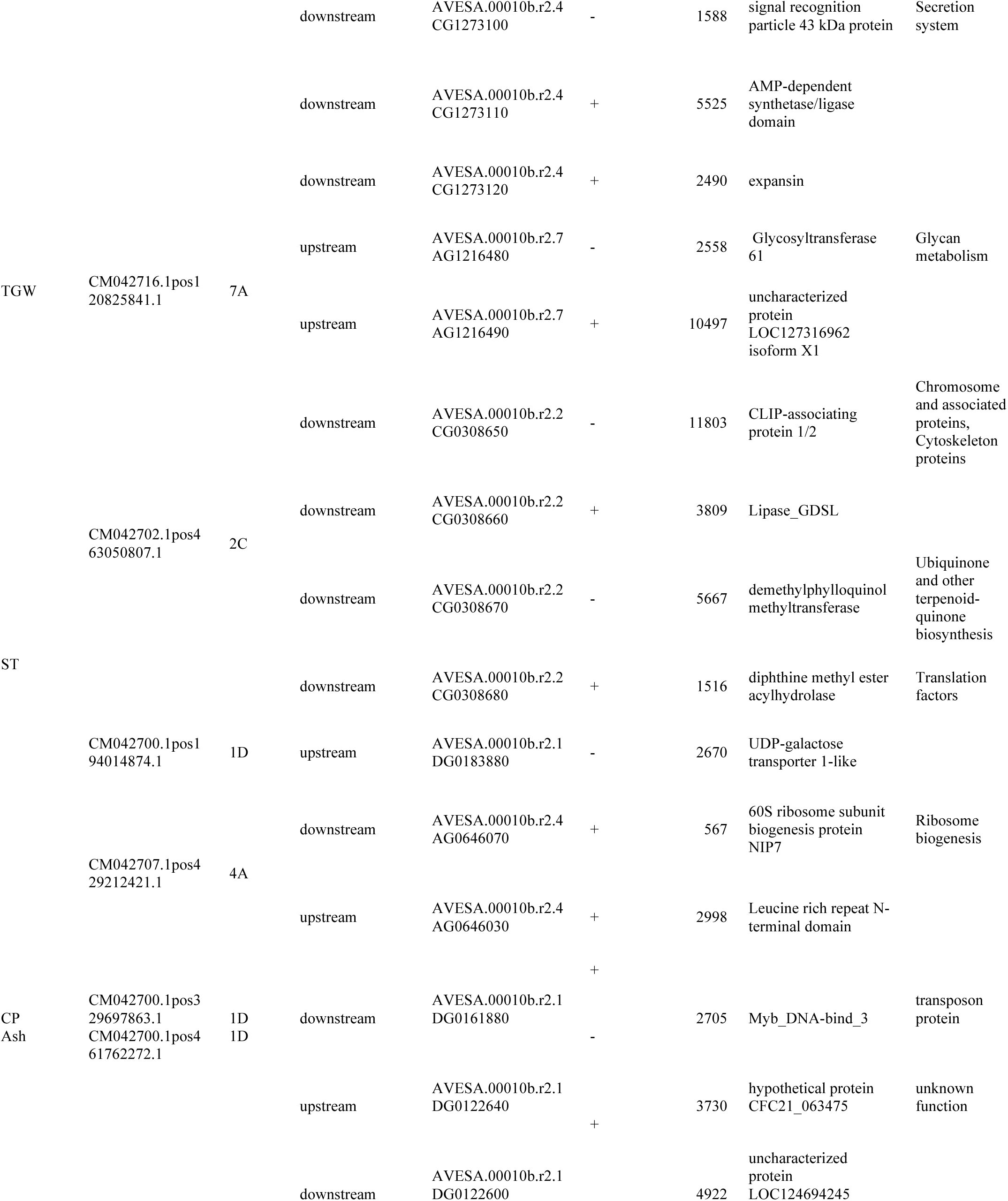
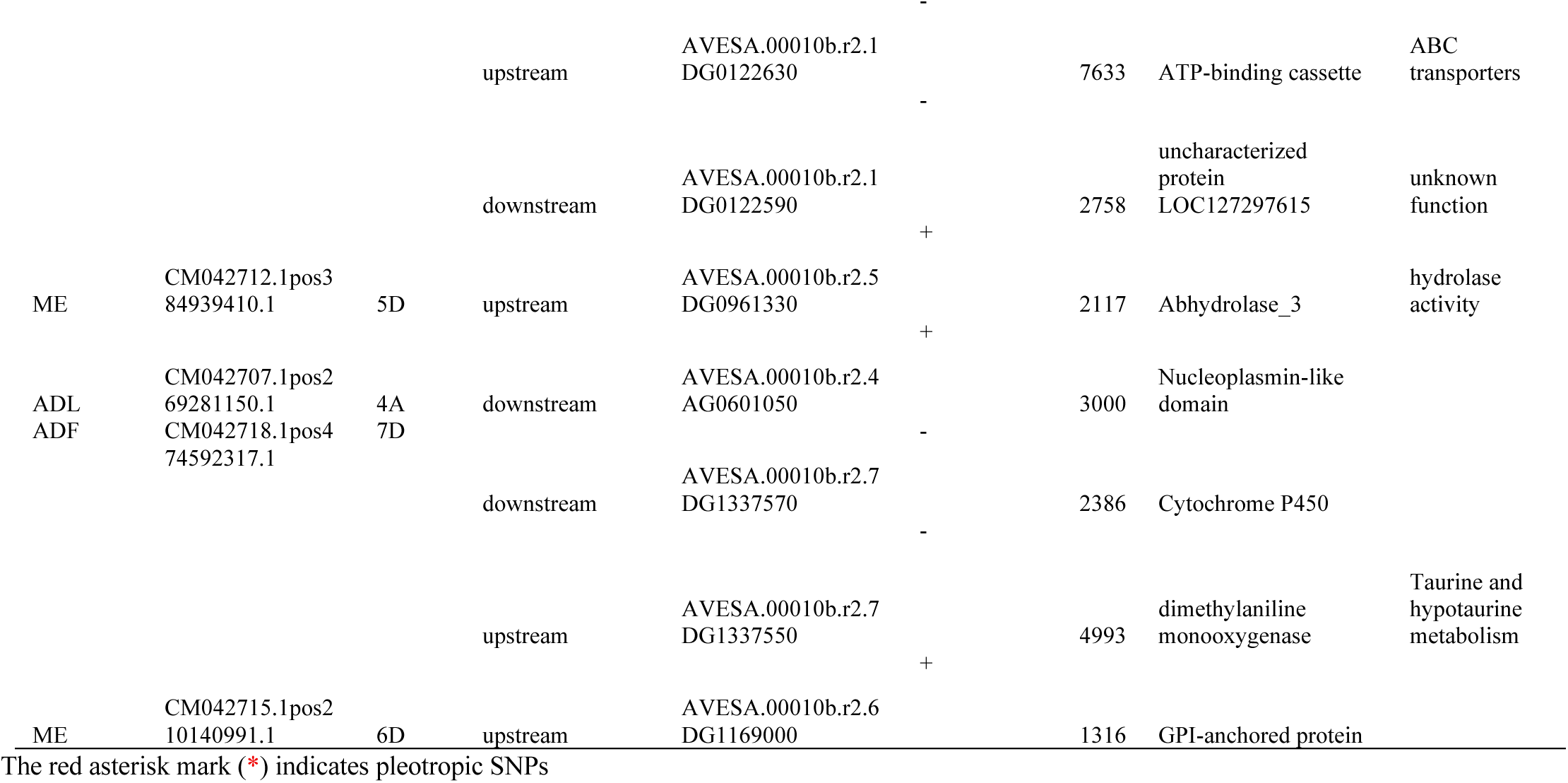
Genomic loci significantly associated with agronomic traits, including chromosomal positions and functional annotations of nearby candidate genes. Gene annotations were obtained using the oat reference database.

## Discussion

### Phenotypic Variation

Analysis of variance revealed significant genetic variation among oat genotypes for all traits, indicating strong potential for selection and genetic improvement. These findings are consistent with previous reports of variability in oat germplasm for similar traits (Krishna et al., 2013; Dubey et al., 2014; Chawla et al., 2022). However, phenotypic variability alone is not sufficient for effective selection; assessing heritability and genetic advance is crucial to determine the portion of variation that is heritable. Traits such as SA and TGW, which showed both high heritability and high genetic advance, are ideal for phenotypic selection, suggesting that these traits are primarily governed by additive gene action and are less influenced by environmental factors, thereby offering greater potential for genetic gain through selection (Johnson et al., 1955; Vimal and Vishwakarma, 1998). In contrast, traits such as ST and ODWT demonstrated both low heritability and low genetic advance, indicating a stronger influence of environmental factors and the predominance of non-additive genetic effects. These traits may require alternative breeding strategies such as genomic selection, and genome editing or environmental management for effective improvement (Duenk et al., 2020). On the other hand, traits like NLPP and plant vigour showed moderate heritability coupled with high genetic advance, indicating moderate potential for improvement. These field evaluations captured substantial genetic variability among the genotypes and responses to variable environmental conditions such as frost, waterlogging, and disease pressure, which is key to identifying resilient lines. These insights are fundamental for advancing promising genotypes through the variety release process and guiding future genetic analyses, including assessments of population structure.

### Genetic diversity and Population Structure

Globally, the number of conserved oats accessions has dropped significantly from 223,287 reported in 1998 to 130,653 by 2010 (FAO, 2010). This downward trend raises concerns for the long-term conservation and accessibility of oat genetic resources. Assessing genetic diversity is therefore crucial, not only to preserving valuable germplasm but also for guiding breeding programs aimed at developing resilient and high-performing varieties.

In this study, 1,823 highly polymorphic SNP markers were used to assess genetic diversity in oat genotypes, with average He and PIC values of 0.42 and 0.33, respectively. These results are slightly higher but generally consistent with those reported by Yan et al. (2020) (He = 0.35, PIC = 0.33) and Wang et al. (2023) (He = 0.32, PIC = 0.26). As standard measures of genetic diversity, the relatively high He and PIC values observed in this study indicate substantial genetic diversity among the oat genotypes. Although we previously noted a global decline in conserved oat accessions, our findings show high genetic diversity within the studied population. This apparent contradiction may be due to factors such as outcrossing, seed exchange among farmers, and gene flow between populations, all of which can enhance within-population diversity even the total number of accessions declines (Khoury et al., 2022; Yirgu et al., 2023). Despite recent studies in major cereals like wheat suggest decline in genetic diversity over time (Sthapit et al., 2020; Gelelcha et al., 2023), such a loss was not observed in oat breeding accessions or cultivars. In contrast, studies in oat have reported an increase in genetic diversity, supporting our findings. For example, Fu et al. (2003) reported increased genetic diversity in oat cultivars released between 1930 and 1950. Similarly, Lyubimova et al. (2020) found a steady increase in average genetic diversity in modern oat breeding varieties from 1929 to 2019. One possible explanation for the higher diversity in oat is the crop’s relatively recent and less intensive breeding history, which has likely resulted in reduced genetic bottlenecking (Yan et al., 2020).

Population structure analysis in this study revealed two distinct genetic groups, consistent with earlier studies, albeit with different panels (Montilla-Bascón et al., 2013; Esvelt Klos et al., 2016; Wang et al., 2023). The presence of admixture in several accessions suggests shared ancestry and gene flow, likely resulting from historical selection, controlled crosses, and occasional natural outcrossing (0.8-1.3%), despite oat being primarily self-pollinated (Yan et al., 2020; Revathi et al., 2023). Earlier studies reported weak population structure in oat (Asoro et al., 2013; Huang et al., 2020), whereas our analysis revealed clearer genetic differentiation. PCA showed the first three components accounted for 29% of total variation, higher than the 20-24% reported previously (Esvelt Klos et al., 2016; Yan et al., 2020) indicating a stronger and more defined genetic structure. The two groups identified by STRUCTURE showed non-overlapping clusters, and the genotype assignments within each cluster were largely consistent with the clusters detected by hierarchical clustering.

### Linkage disequilibrium (LD) Analysis

Linkage disequilibrium (LD) analysis revealed numerous significant marker pairs, with a mean r² of 0.25 (Supplementary Table 9). Since r² values above 0.2 are effective for detecting causal loci, particularly in centromeric regions (Alqudah et al., 2020), most markers used in this study are well-suited for association mapping. Genome-wide LD decayed at 7.4 Mb, lower than the 2.29 Mb reported in a global oat panel (Peng et al., 2022), but faster than the 28 Mb observed in advanced breeding lines (Bazzer et al., 2025), indicating moderate GWAS resolution in this panel. As a predominantly self-pollinated species, oat typically shows slower LD decay, allowing for effective genome coverage with fewer markers (Flint-Garcia, 2003; Alqudah et al., 2020). LD decay informs QTL confidence intervals and guides marker density requirements (Sallam & Martsch, 2015; Otyama et al., 2019), with previous studies recommending at least one marker per LD decay unit (Yan et al., 2020; Wang et al., 2023). Given oat’s estimated 10 Gb genome size (Peng et al., 2022), approximately 1,500 markers would suffice, for GWAS analysis, yet the 18,196 high-quality SNPs used here far exceed this benchmark, enhancing the resolution and robustness of our association mapping.

### Marker Trait Associations for forage-related traits

A total of 42 significant SNPs associated with sixteen agronomic traits were identified, with 1 to 8 markers detected per trait (FDR *p* ≤ 0.05, Bonferroni threshold = 0.15). These SNPs were distributed across 13 distinct chromosomes of the A, C, and D subgenomes, with the highest density on chromosome 5C (10 SNPs). Most SNPs were specific to environments, with only three (CM042711.1pos74917192.1 associated with SW, CM042705.1pos607673018.1 associated with ADL and ME, and CM042700.1pos329697863.1 associated with CP) showing consistency across multiple sites, likely reflecting the strong environmental influence and complex polygenic nature of these traits (Rispail et al., 2018).

Numerous pleiotropic loci have been reported in previous GWAS across various crops, including rice and wheat (Ashfaq et al., 2023; Zhao et al., 2024). Likewise, this study identified four pleiotropic SNPs associated with grain-related and forage quality traits, suggesting interdependence or pleiotropic genetic loci (Table 3). These loci offer strong potential for QTL pyramiding and marker-assisted selection to improve multiple grain traits simultaneously in oat. Previous association studies in oats were limited by the lack of a reference genome, relying instead on consensus maps and reporting QTLs in centimorgans (cM). The recent release of a high-quality oat reference genome (Peng et al., 2022), enabled this study to anchor GBS-derived SNPs to physical positions, significantly improving mapping resolution and reducing positional ambiguity. Although direct comparisons with earlier cM-based QTLs remain challenging, the genomic framework now enables more precise localization of trait-associated loci.

### Candidate Gene Identification and Functional Annotation

This study identified 42 candidate genes near SNPs significantly associated with key traits, particularly seed-related characteristics (seed length, width, area, and thousand-grain weight) as well as with other agronomic traits (plant height, leaf length, and stem thickness) and feed quality traits (ADL, ME and IVOMD). One notable gene, AVESA.00010b.r2.4CG1272870.1, encodes adenylate kinase, homolog to *OsAK3* in rice, which regulates grain length in close proximity to CM042708.1pos82518881.1 on chromosome 4C SNP. Mutant of *OsAK3* produce shorter grains, while overexpression increases grain size (Zhang *et al.,* 2021). Another candidate gene, AVESA.00010b.r2.4CG1272900, near to same SNP (CM042708.1pos82518881.1) annotated as *UDP-glucosyltransferase*, aligns with rice gene *GSA1*, where overexpression enhances grain size (Dong et al., 2020). These findings highlight the critical trade-off between oat grain sizes and forage quality, particularly in dual-purpose varieties (Shah et al., 2020; Ertekin et al., 2023). Larger-grain oats typically offer high grain yield and later maturity but often compromise forage quality due to increased stem content and lignification when harvested late (Chapko et al., 1991). In contrast, smaller-grain or forage-type oats provide greater vegetative biomass, higher crude protein, and improved digestibility when harvested at the boot stage (Liu and Mahmood., 2015). Understanding this balance is essential for optimizing both grain and forage yield in integrated crop-livestock systems and the SNP marker identified here could facilitate marker assisted breeding. Another transcript, AVESA.00010b.r2.4AG0646010, located near a pleiotropic SNP (CM042707.1pos429035978.1) associated with multiple traits including plant height and leaf length, encodes zeta-carotene isomerase (Z-ISO), a key enzyme in carotenoid biosynthesis (Efremov et al., 2021). A study conducted in rice revealed that mutations in the Z-ISO gene result in high-tillering and dwarf phenotypes (Liu et al., 2021), highlighting the gene’s contribution to these traits. Since plant height is a key trait contributing to forage biomass yield in oat (Tessema & Getinet, 2020), this finding provides valuable insight for future oat breeding programs targeting forage improvement.

In the vicinity of significant SNP CM042711.1pos74917192.1 on chromosome 5C a promising candidate gene, AVESA.00010b.r2.5CG0924570, encoding a subunit of the endosomal sorting complex required for transport, II (*ESCRT-II*), identified. This gene plays a critical role in intracellular trafficking, signal transduction, and cellular development. Its rice homolog *OsVPS22* has been shown to influence grain quality by regulating chalky endosperm formation and affecting seedling viability, highlighting its essential role in developmental processes (Zhang et al., 2013). In forage crops, homologous genes in the ESCRT pathway have analogous functions such as regulating tissue differentiation, cell wall composition and carbohydrate partitioning, which are directly linked to digestibility, biomass quality, and early plant vigor (Jung and Allen 1995; Hatfield et al., 2007; Bosch et al 2011).

Another transcript, AVESA.00010b.r2.7DG1337570, located near a SNP associated with the feed quality trait ADL, encodes a cytochrome P450 enzyme. Cytochrome P450s are well known for their role in the phenylpropanoid pathway, particularly in lignin biosynthesis. A study in rice confirmed that this enzyme contributes to lignin biosynthesis (Supatmi et al., 2025). Similarly, a study in barley reported that mutations in a cytochrome P450 gene caused cutin layer instability (Ameen et al., 2021). Together, these findings suggest that the association between the identified SNP and this candidate gene is biologically plausible and warrants further investigation.

## Conclusion

The oat accessions analyzed in this study displayed substantial genetic diversity, highlighting significant potential for improving oat production and productivity in Ethiopia. This diversity serves as a valuable resource for developing new oat varieties with enhanced traits such as higher yield, improved environmental adaptability, and better feed and food qualities. GWAS identified 42 SNPs significantly associated with sixteen traits, including feed quality, vegetative traits, and grain size–related traits, along with 46 candidate genes near these loci. These candidate genes span diverse families such as enzymes, F-box proteins, regulatory factors, membrane trafficking components, and structural proteins. Their identification provides valuable insights into the genetic basis of traits analyzed and offers markers for targeted breeding. Integrating these candidate genes and SNPs into breeding programs will accelerate the development of high-yielding, climate-resilient oat varieties, supporting food security and agricultural productivity under changing environmental conditions.

## Acknowledgment

The authors would like to thank Dr Elena Grosu, for reading and commenting on the original manuscript. We also would like to thank Mr. Gezahegn Mengistu (EIAR) and Ms. Samrawit Fesseha (Addis Ababa University) for technical support during the field trials. We also wish to acknowledge Mr. Yonas Asmare for his valuable technical support during the feed quality analyses.

## Data availability statement

The data generated and presented in this study is in the process of being deposited in a public database.

## Author contribution

TA, JDG, AL and BS designed and supervised the project and the manuscript writing, TA, JDG, AL and CJ analyzed phenotypic and genotypic datasets. AL and LH collected leaf samples and extracted DNA. DM, MK, FF, LH and NTA involved in the supervision of the phenotyping and feed quality analysis. MT, RA, TA and BS and JDG involved in the supervision of the genotyping project. All authors have read, reviewed and approved this manuscript.

## Declarations

Ethics Approval and Consent to Participate Not applicable.

## Consent for Publication

Not applicable.

## Supplementary Tables

**Sup. Table 1.** Metadata for 169 oats accessions used in this study

**Sup. Table 2.** Traits measured in this study along with the measurement methods employed

**Sup. Table 3.** Combined analysis of variance (ANOVA) for agro-morphological, feed quality, and grain-related traits of oat accessions evaluated across three locations over two years.

**Sup. Table 4.** Principal component loadings and explained variance for 21 traits in 167 oat genotypes

**Sub. Table 5.** Clusters grouped by STRUCTURE software for 167oat accessions at K = 2

**Sub. Table 6.** Clusters grouped by Neighbor Joining method for 167 oat accessions

**Sub. Table 7.** Analysis of molecular variance (AMOVA) results among and within populations

**Sub. Table 8.** Summary of linkage disequilibrium analyses among marker pairs and the number of significant marker pairs per chromosome and genome

**Sub. Table 9.** Genome-wide association results for key agronomic traits showing chromosome position, p-value, SNP, minor allele frequency (MAF), effect and model results for Marker-Trait Associations for selected agronomic traits.

